# The Entorhinal-DG/CA3 Pathway in the Medial Temporal Lobe Retains Visual Working Memory of a Simple Surface Feature

**DOI:** 10.1101/2022.08.31.506098

**Authors:** Weizhen Xie, Marcus Cappiello, Michael A. Yassa, Edward Ester, Kareem Zaghloul, Weiwei Zhang

## Abstract

Classic models consider working memory (WM) and long-term memory as distinct mental faculties that are supported by different neural mechanisms. Yet, there are significant parallels in the computation that both types of memory require. For instance, the representation of precise item-specific memory requires the separation of overlapping neural representations of similar information. This computation has been referred to as pattern separation, which can be mediated by the entorhinal-DG/CA3 pathway of the medial temporal lobe (MTL) in service of long-term episodic memory. However, although recent evidence has suggested that the MTL is involved in WM, the extent to which the entorhinal-DG/CA3 pathway supports precise item-specific WM has remained elusive. Here, we combine an established orientation WM task with high-resolution fMRI to test the hypothesis that the entorhinal-DG/CA3 pathway retains visual WM of a simple surface feature. Participants were retrospectively cued to retain one of the two studied orientation gratings during a brief delay period and then tried to reproduce the cued orientation as precisely as possible. By modeling the delay-period activity to reconstruct the retained WM content, we found that the anterior-lateral entorhinal cortex (aLEC) and the hippocampal DG/CA3 subfield both contain item-specific WM information that is associated with subsequent recall fidelity. Together, these results highlight the contribution of MTL circuitry to item-specific WM representation.

## Introduction

Working memory (WM) actively retains a small amount of information to support ongoing mental processes ^1^. This core mental faculty relies upon distributed brain regions ^2^, ranging from lower-level sensory areas ^3^ (but see ^4^) to higher-level frontoparietal networks ^4–7^. This distributed neocortical network, however, often does not involve the medial temporal lobe (MTL), which is traditionally attributed to long-term episodic memory ^9,10^. This distinction is grounded in the separation between WM and long-term memory in classic models ^11,12^ and in early MTL lesion case studies ^13,14^. Yet, this classic view is not free of controversy. A growing body of research has suggested that the MTL is involved in tasks that rely on information maintained in WM ^15–24^. Furthermore, MTL lesions can disrupt WM task performance ^25–28^. Despite these recent findings, however, major theories have not considered the MTL as a mechanism underlying WM representation ^8,29^. First, it is unclear what computation process of the MTL is involved in WM ^29^. Furthermore, the MTL tends to engage more in a WM task when long-term memory becomes relevant, for example when task loads are higher ^17,21,23^ or when task stimuli are complex ^15,19,20,24,30,31^. As a result, contributions from the MTL to WM are often deemed secondary ^8,29^.

Clarifying this issue requires specifying how the MTL contributes to WM representation and the extent to which this contribution holds even when WM task demand is minimized. Although WM and long-term memory are traditionally considered separate mental faculties, the functional parallels in both types of memory suggest potential shared neural mechanisms ^32–35^. For example, the ability to retain precise item-specific memory would require the computation to distinguish neural representations of similar information – a process known as pattern separation ^36^. This aspect of long-term memory is widely thought to emerge from various properties of the MTL’s entorhinal-DG/CA3 pathway ^36–43^, such as abundant granule cells and strong inhibitory interneurons in the hippocampal DG, as well as powerful mossy fiber synapses between the DG and CA3 subfields ^42,44^. These properties make it possible to enable sparse coding to ensure a sufficient representational distance among similar information ^45,46^. As these hippocampal substructures communicate with other neocortical areas via the entorhinal cortex ^38,42^, there is a proposed gradian of pattern separation along the entorhinal-DG/CA3 pathway to support item-specific long-term episodic memory ^37^. These ideas are supported by evidence based on animal and human behaviors ^47–49^, electrophysiological recordings ^50–52^, and human fMRI ^37,38,40,53^. However, the extent to which the entorhinal-DG/CA3 pathway is involved in WM, especially in humans other than animal models ^54^, has remained unclear.

Several challenges faced in past research may add to this uncertainty. For example, it is difficult to infer signals from MTL substructures, especially those within the hippocampus, based on human fMRI using a standard spatial resolution ^4,5^ or intracranial direct recording with limited electrode coverage ^19–22^. Furthermore, the use of complex task designs with multiple memory items ^31^ might also be suboptimal to reveal item-specific WM information in MTL subregions without taxing too much on the WM storage limit. To investigate these issues, here, we leverage an established retro-cue orientation WM task ^3–5^ and a high-resolution fMRI protocol to test the key prediction that the MTL’s entorhinal-DG/CA3 pathway retains item-specific WM information of a simple surface feature. In this task, participants are directed to retain the orientation information of a cued stimulus from two sequentially presented orientation gratings (separated at least by 20°; **Figure 1A**). After a short delay (5 TRs; 1TR = 1.75s), they try to reproduce the cued orientation grating as precisely as possible using the method of adjustment. As participants are retrospectively cued to retain only one item during the delay, they are expected to encode both items but then only keep one in mind during the delay period. This design imposes a task demand on the observer to correctly select the cued orientation while resisting the interference from the internal representations of other similar orientation gratings. This post-encoding information selection function during a short delay has been considered a hallmark of WM ^55,56^, regardless of the presence or absence of sustained neural activation ^57,58^. If the MTL’s entorhinal-DG/CA3 pathway indeed supports this function, it is expected that the recorded delay-period activity should contain more information about the cued item, as compared with the uncued item, even though both items are initially remembered with an equal likelihood ^3–5^. If, however, information about the cued item and the uncued item is equally present during the delay period, the MTL may play a limited role in the representation of task-relevant information in WM but more during the initial encoding.

**Figure 1.**
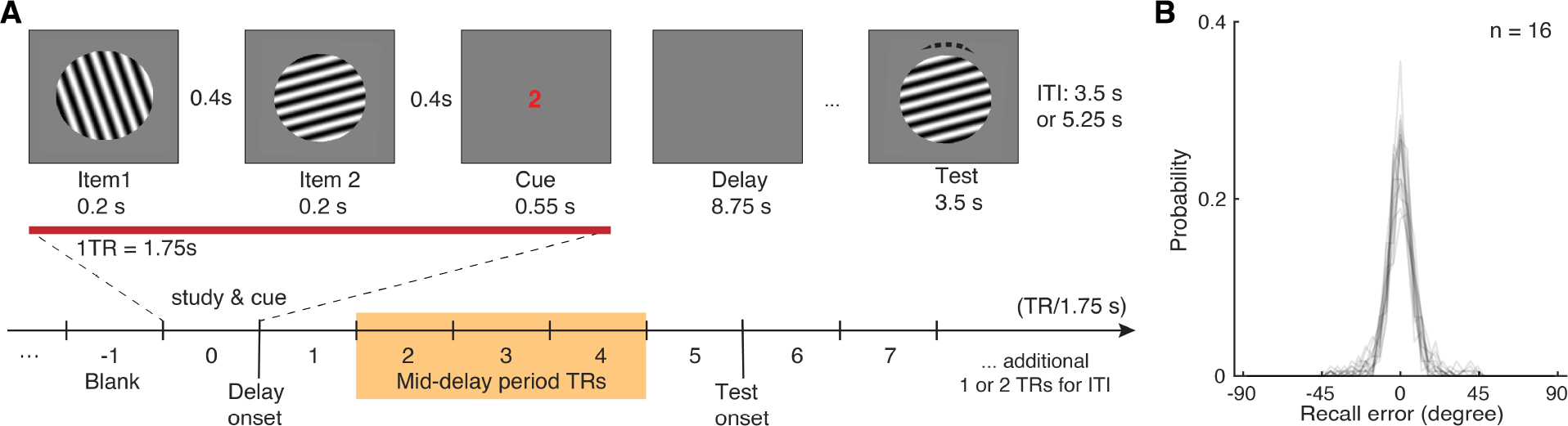
Visual WM task and participants’ task performance. (**A**) During fMRI scanning, participants were directed to retain the orientation of a cued grating stimulus from two sequentially presented grating stimuli (item 1 vs. 2). After a short retention interval, they tried to reproduce the cued orientation grating as precisely as possible. (**B**) Participants’ task performance was high and mostly driven by the fidelity of the retained visual WM content.Each trace represents a participant’s recall probability in the feature space (−90 to 90 degrees) separated by 45 bins. TR = MR repetition time; ITI = inter-trial interval. The shaded area in (**A)** highlights the middle 3 TRs of the delay period.

## Results

Participants’ memory performance is quantified as recall error – the angular difference between the reported and the actual orientations of the cued item ^59^. As the effective memory set size is low at one memory item, participants’ performance is high with an average absolute recall error of 12.01° ± 0.61° (mean ± s.e.m.). Furthermore, the recall error distribution is centered around 0° with most absolute recall errors smaller than 45° (~97% trials; **Figure 1B**). These behavioral data suggest that participants in general have remembered high-fidelity orientation information of the cued item during the delay period.

### Fine discrimination of Remembered WM Content in the MTL

Of primary interest, we examined whether precise orientation information of the cued item is retained during WM retention in anatomically-defined MTL regions of interest (ROIs; **Figure 2A**), including the entorhinal cortex (anterior-lateral, aLEC & posterior-medial, pMEC), the perirhinal cortex, para-hippocampus, and hippocampal DG/CA3, CA1, subiculum, as defined in the previous studies ^53,60^. Additionally, we chose the amygdala as a theoretically irrelevant but adjacent control region, because the involvement of the amygdala for emotionally neutral orientation information is expected to be minimal ^61^. This allows us to gauge the observations in MTL ROIs while controlling for the signal-to-noise ratio in fMRI blood-oxygenation-level-dependent (BOLD) signals in deep brain structures.

**Figure 2.**
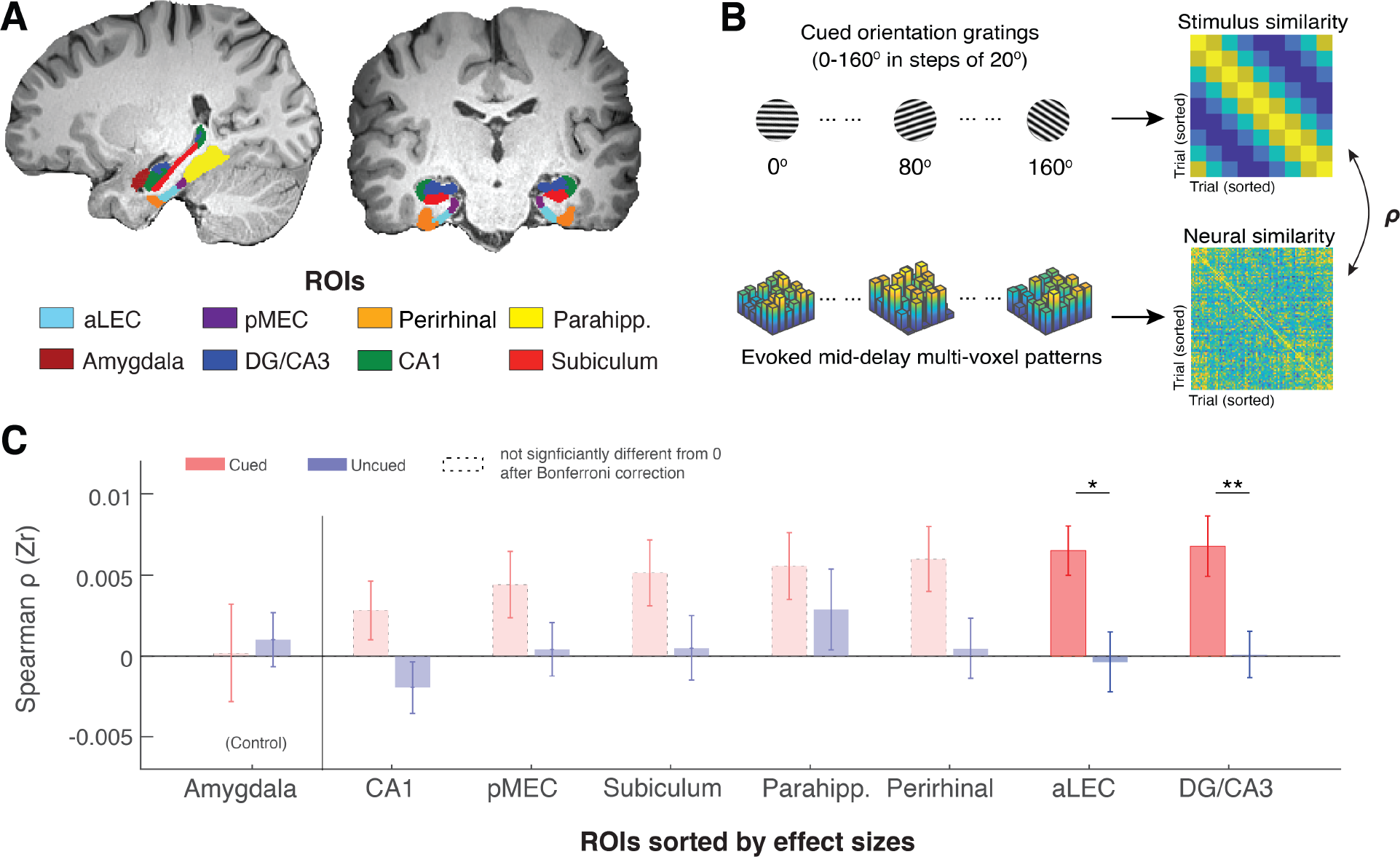
The MTL retains item-specific WM information revealed by stimulus-based representational similarity analysis. (**A**) MTL ROIs are parcellated based on previous research ^53,60^. The amygdala is chosen as an adjacent control region. (**B**) For each ROI, we examined the extent to which the evoked multi-voxel pattern during the mid-delay period could keep track of the feature values among different WM items. Specifically, we correlated the feature similarity of every two cued items with the similarity in their evoked neural patterns during the WM delay period. If a brain region contains item-specific information to allow fine discrimination of different items, the evoked neural patterns should keep track of the feature similarity of these items ^63^. (**C**). Across ROIs, we find that this prediction is supported by data from the aLEC and DG/CA3, which show a larger effect size in the association between neural similarity and stimulus similarity based on the cued item as compared with the uncued item. Error bars represent the standard error of the mean (s.e.m.) across participants. \**p* < 0.05 and \*\**p* < 0.01 for the comparison of the results based on cued versus uncued items; aLEC = anterior-lateral entorhinal cortex; pMEC = posterior-medial entorhinal cortex; parahipp. = parahippocampus. Results from detailed statistical tests are summarized in **Table S1**.

As recent neural theories of WM have proposed that information retained in WM may not rely on sustained neural activation ^5,58,62^, we inspected how the multivoxel activity pattern in each subject-specific ROI is correlated with the retained WM content predicted by the cued orientation gating (**Figure 2B**). We found that certain voxels in an ROI could respond more strongly to a particular cued orientation, even when the average BOLD activity across voxels does not show preferred coding for a certain orientation (see an example in **Figure S1**). We then assessed the consistency of these stimulus-related multivoxel activity patterns in the MTL and the amygdala control region based on stimulus-based representational similarity analysis. In this analysis, we correlated the angular similarity of every pair of cued orientation gratings with the similarity of the evoked BOLD patterns in these trials. The rationale is that if orientation information is retained within an ROI, the recorded neural data should track the relative angular distance between any two cued orientation gratings (hence fine discrimination ^63^). Informed by the previous research ^3,5^, we performed this analysis using the raw fMRI BOLD signals from the middle 3TRs out of the 5-TR retention interval to minimize the contribution of sensory process or anticipated retrieval, hence maximizing the inclusion of neural correlates of WM retention ^64^. A time-varying version of this analysis is summarized in **Figure S6**.

In line with our prediction, we found that stimulus similarity for the cued item was significantly correlated with neural similarity across trials in the aLEC (t(15) = 4.29, p = 6.48e-04, p_Bonferroni_ = 0.0052, Cohen’s d = 1.11) and DG/CA3 (t(15) = 3.64, p = 0.0024, p_Bonferroni_ = 0.019, Cohen’s d = 0.94; **Figure 2C**). In contrast, stimulus similarity for the uncued item across trials could not predict these neural similarity patterns (p’s > .10). Furthermore, the evoked neural similarity patterns in these regions were significantly more correlated with the cued item than with the uncued item (aLEC: t(15) = 2.66, p = 0.018, Cohen’s d = 0.69; DG/CA3: t(15) = 3.64, p = 0.0024, Cohen’s d = 0.94). While the rest of the MTL showed similar patterns, we did not obtain significant evidence in other MTL ROIs following the correction of multiple comparisons (see **Table S1** for full statistics), suggesting attenuated effect sizes in these regions. Furthermore, neural evidence related to the cued item in the aLEC and DG/CA3 was significantly stronger than that in the amygdala control ROI. This was supported by a significant cue (cued vs. uncued) by region (combined aLEC-DG/CA3 vs. amygdala) interaction effect on the correlation between stimulus and neural similarity patterns (F(1, 15) = 4.97, p = 0.042). Together, these results suggest that delay-period activity patterns in the entorhinal-DG/CA3 pathway are associated with retrospectively selected task-relevant information, implying the presence of item-specific WM representation in these subregions.

### Reconstruction of Item-specific WM Information based on Inverted Encoding Modeling

To directly reveal the item-specific WM content, we next modeled the multivoxel patterns in subject-specific ROIs using an established inverted encoding modeling (IEM) method ^5^. This method assumes that the multivoxel pattern in each ROI can be considered as a weighted summation of a set of orientation information channels (**Figure 3A**). By using partial data to train the weights of the orientation information channels and applying these weights to an independent hold-out test set, we reconstruct the assumed orientation information channels to infer item-specific information for the remembered item – operationalized the resultant vector length of the reconstructed orientation information channel normalized at 0° reconstruction error (**Figure S2**). As this approach verifies the assumed information content based on observed neural data, its results are interpretable within the assumed model, even though the underlying neuronal tuning properties are unknown ^5,65^. Based on this method, previous research has revealed item-specific WM information in distributed neocortical areas, including the parietal, frontal, and occipital-temporal areas ^4,5,66,67^. We have replicated these effects in the current dataset (**Figure S3)**.

**Figure 3.**
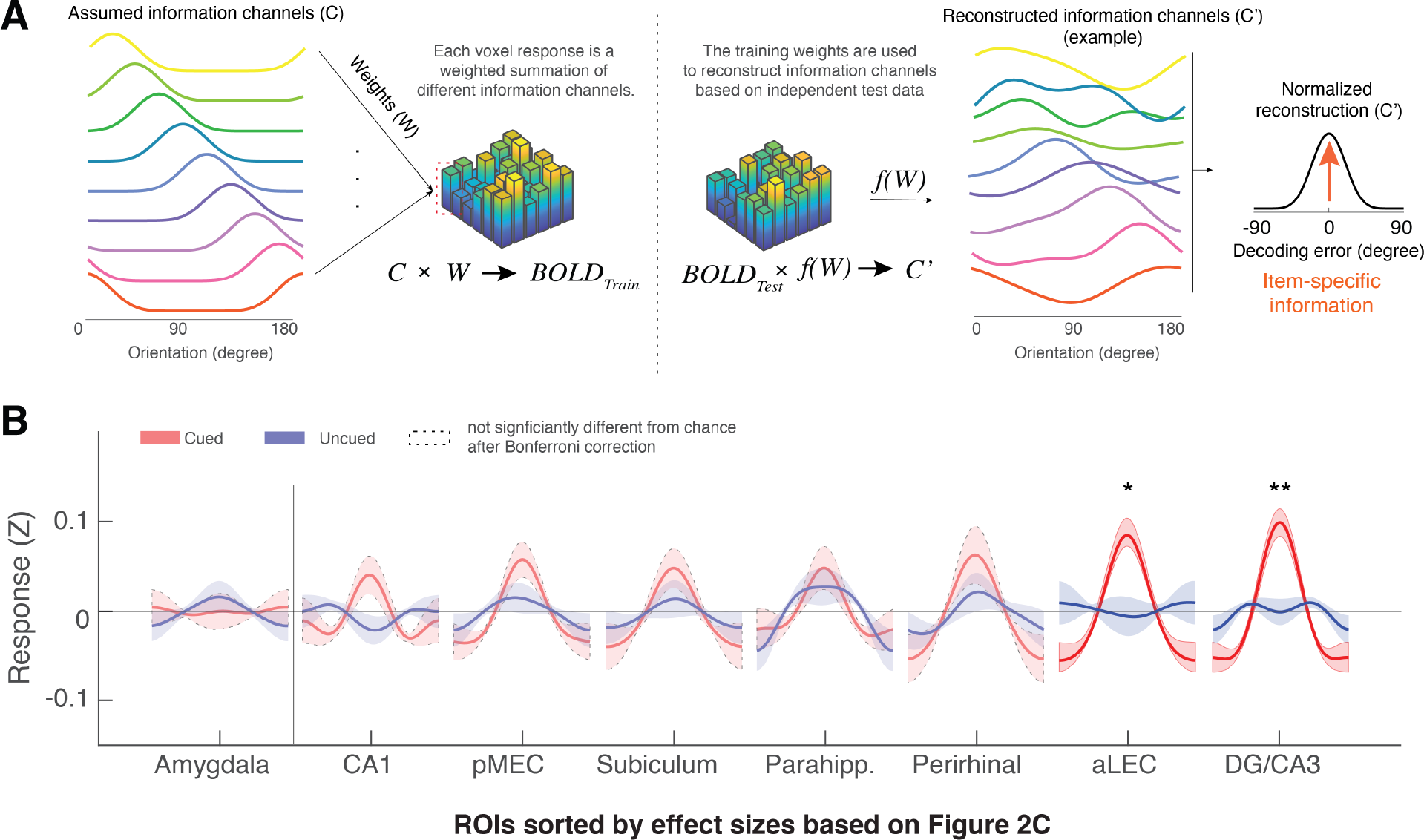
The MTL retains item-specific WM information revealed by Inverted Encoding Modeling (IEM). (**A**) The IEM method assumes that each voxel response in the multi-voxel pattern reflects a weighted summation of different ideal stimulus information channels (*C*). The weights (*W*) of these information channels are learned from training data and then applied to independent hold-out test data to reconstruct information channels (*C’*). After shifting these reconstructed information channels to a common center, the resultant vector length of this normalized channel response reflects the amount of retained information on average (also see **Figure S2**). (**B**) We find that the BOLD signals from both the aLEC and DG/CA3 contain a significant amount of item-specific information for the cued item, relative to the uncued item. Shaded areas represent the standard error of the mean (s.e.m.) across participants. To retain consistency, we sorted the *x*-axis (ROIs) based on **Figure 2C**. \**p* < 0.05 and \*\**p* < 0.01 for the comparison of the results based on cued versus uncued items; a.u. = arbitrary unit; aLEC = anterior-lateral entorhinal cortex; pMEC = posterior-medial entorhinal cortex; parahipp. = parahippocampus. Results from detailed statistical tests are summarized in **Table S2**.

Moving beyond these well-established observations in distributed neocortical structures, we found that the amount of reconstructed item-specific information for the cued item during WM retention was also significantly greater than chance level in two anatomically defined MTL subregions, aLEC (t(15) = 4.41, p = 5.07e-04, p_bonferroni_ = 0.0041, Cohen’s d = 1.14) and the hippocampal DG/CA3 (t(15) = 4.73, p = 2.68e-04, p_bonferroni_ = 0.0021, Cohen’s d = 1.22; **Figure 3B**). These effects were specific to the maintenance of the cued item, as information related to the uncued item was not statistically different from chance (p’s > .10) and was significantly less than that for the cued item (aLEC: t(15) = 2.75, p = 0.015, Cohen’s d = 0.71; DG/CA3: t(15) = 3.83, p = 0.0016, Cohen’s d = 0.99). Critically, the amount of information specific to the cued item in the aLEC and DG/CA3 was significantly greater than that in the amygdala control ROI, which is supported by a significant cue (cued vs. uncued) by region (combined aLEC-DG/CA3 vs. amygdala) interaction effect on IEM reconstruction outcomes (F(1, 15) = 7.16, p = 0.016).

Collectively, results from complementary analytical procedures suggest that the MTL’s entorhinal-DG/CA3 pathway retains precise item-specific WM content for a simple surface feature (e.g., orientation) to allow fine discrimination of different items in the feature space. As such, the stimulus-based prediction of neural similarity is highly correlated with the amount of reconstructed information based on IEM, even though these two analyses are based on different analytical assumptions (e.g., correlation between IEM and representational similarity analysis for the cued item, aLEC: r = 0.87, p < .0001; DC/CA3: r = 0.78, p < .0001, **Figure S4**).

### Reconstruction of WM Item Information in the MTL is associated with recall fidelity

Next, we examined the extent to which WM information retained in the MTL’s aLEC-DG/CA3 circuitry is related to an observer’s subsequent recall behavior. As the angular resolution of the reconstructed orientation information is 20° in the current study, our data therefore suggest that the MTL can distinguish similar orientation information in WM that is at least 20° apart. This neural separation should be consequential for later recall performance, in that trials with greater item-specific information reconstructed from the MTL should be associated with higher WM recall fidelity. To test this prediction, we grouped the trials from each participant into two categories. The first category contains small recall error trials, where participants make an effective recall response within one similar item away from the cued item (absolute recall error < 20°; 149 ± 3 trials [mean ± s.e.m.]). Another category contains larger recall error trials (27 ± 3 trials) with absolute recall errors that are greater than 20° but smaller than the 3 standard deviations (SD) of the aggregated recall error distribution (**Figure 4A**). These trials would capture participants’ imprecise recall responses for the cued item, instead of those with an extra-large recall error that could be attributed to other factors such as attentional lapses ^68^. The two identified categories of trials together account for about 98% of the total trials (i.e., 176 out of 180 trials). We have obtained similar results based on another thresholding heuristic by just using 45° of absolute recall error as a cut-off (i.e., half of the 90° range; see **Supplementary Information**).

**Figure 4.**
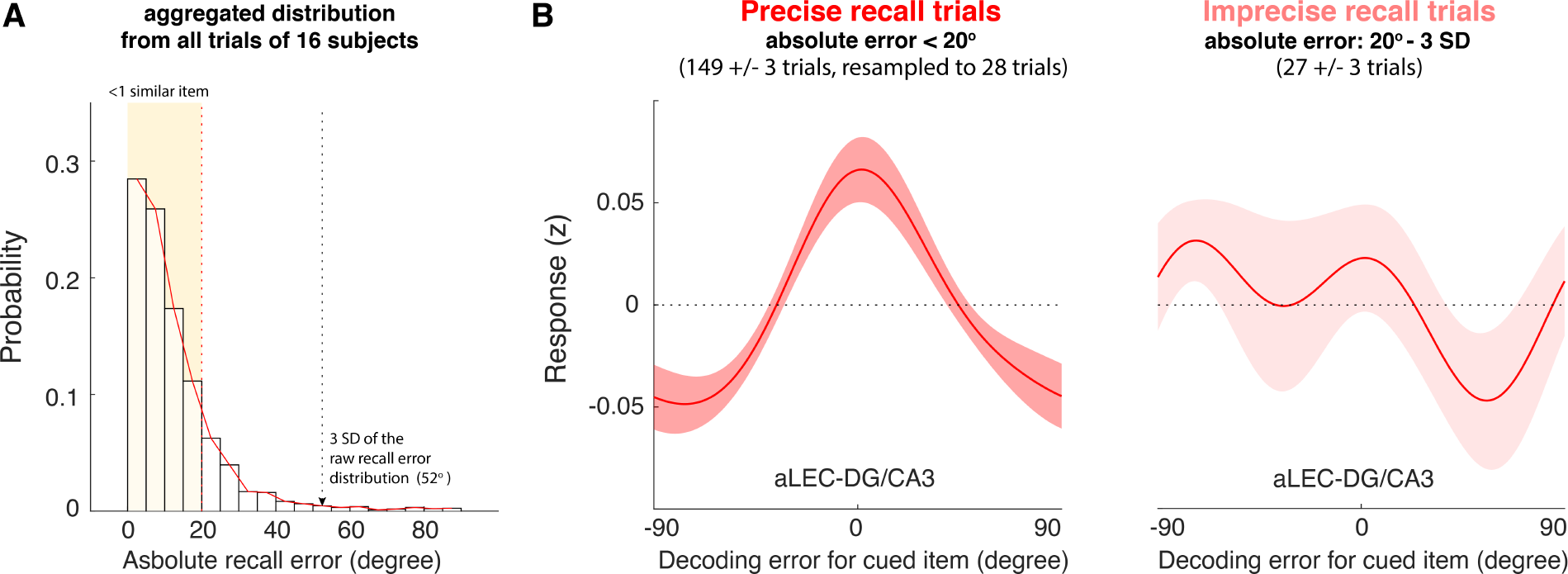
The quality of WM information retained in the aLEC-DG/CA3 pathway is associated with later recall fidelity. (**A**) Participants’ performance in the visual WM task was high with about 98% of absolute recall errors falling within the 3 SD of the aggregated recall error distribution. As the angular resolution of the presented orientation grating is at least 20° between any two items, for most of the trials, participants’ recall responses were as precise as within one similar item away from the cued item (absolute recall error < 20°). (**B**) By inspecting the IEM reconstructions for trials with small errors (absolute error < 20°) and trials with larger errors (absolute recall error: 20° to 3 SD of recall errors), we find that the quality of IEM reconstructions in the combined aLEC-DG/CA3 ROI varies as a function of participants’ recall fidelity. Precise recall trials have yielded better IEM reconstruction quality, even after resampling the same number of trials from the data to control for imbalanced trial counts between small- and larger-error trials. Shaded areas represent the standard error of the mean (s.e.m.) across participants.

We performed the leave-one-block-out analysis to obtain trial-by-trial IEM reconstructions based on delay-period BOLD signals aggregated from the aLEC and DG/CA3. We then averaged the IEM reconstructions from the small- and larger-error trials separately. As the trial counts between categories were not balanced, we resampled the data from the small-error trials based on the number of larger-error trials for 5,000 times. We took the average of IEM reconstruction across iterations to obtain robust subject-level trial-average estimates with a balanced trial count across different behavioral trial types ^24,69,70^. By contrasting these estimates at the subject level, we found that the small-error trials yielded significant IEM reconstructions for the cued item (t(15) = 4.50, p = 4.21e-04, Cohen’s d = 1.16), whereas the larger-error trials did not (t(15) = 0.03, p = 0.98, Cohen’s d = 0.007, **Figure 4B**). Furthermore, the reconstructed WM information in the combined aLEC-DG/CA3 showed better quality in the small-error trials, as compared with that in the larger-error trials (t(15) = 2.45, p = 0.027, Cohen’s d = 0.61). These results suggest that higher-quality WM representation in the entorhinal-DG/CA3 pathway during the delay period is associated with better subsequent recall fidelity.

## Discussion

Based on high-resolution fMRI, this current study uncovers an often-neglected role of the MTL’s the entorhinal-DG/CA3 pathway in item-specific WM representation at a minimal task load. Our data suggest that the entorhinal-DG/CA3 circuitry retains item-specific information to allow fine discrimination of similar WM items across trials. The quality of item-specific WM information in the entorhinal-DG/CA3 pathway is associated with an observer’s subsequent recall fidelity. Together, these findings fill a missing link in the growing literature regarding how the MTL contributes to WM ^22,29^.

Theoretically, our findings align well with the established literature on the entorhinal-DG/CA3 circuitry and the formation of high-fidelity long-term episodic memory ^36–43^. This function has been linked with various neuronal properties along the entorhinal-DG/CA3 pathway – such as abundant granule cells, strong inhibitory interneurons, and powerful mossy fiber synapses – which could enable sparse coding of information to minimize mnemonic interference ^42,44–46^. As such, similar information can be retained with a sufficient representational distance to support behavioral discrimination ^37,38,40,47–53^. Our data add to this literature by supporting a parsimonious hypothesis that the same MTL mechanism can also be used to support the quality of WM representation ^71^. Specifically, potential interference between items either across or within trials would place a demand on pattern separation even over a short delay ^72^. As such, the MTL circuitry involved in the resolution of mnemonic interference ^42^ would play a key role in reducing inference between WM content and other similar information in the feature space. These findings, therefore, clarify the functional or computational relevance of the MTL in WM, in contrast to the classic view that the MTL is only secondary for WM ^8,29^. These interpretations are consistent with the recent development of neural theories of WM that have highlighted the involvement of distributed brain areas ^2,29,73^, including mechanisms in the MTL that are traditionally deemed irrelevant for human WM ^25,31,32,74^.

Empirically, our results have resolved an issue concerning the decodability of item-specific WM content in the MTL for simple stimulus features. Previously, MTL activity has been shown to scale with WM set size of letters and color squares without decodable item-specific WM content ^21,23^. One conceptual uncertainty is whether the MTL primarily responds to task difficulty or retains item-level information in WM. Our data suggest that the MTL retains item-level WM information even when the effective WM set size is one. The lack of significant observations in some previous studies using the same paradigm may be due to the lack of granularity in MTL recordings ^5^. To investigate this, we aggregated data from all the voxels in the hippocampus to examine whether blurred MTL signals would be sufficient to reveal item-specific WM content using the current IEM procedure. As CA1 and subiculum voxels contain less robust WM information (**Figure 3B**), we predicted that this aggregation procedure would attenuate the evidence for WM information due to the reduction in signal-to-noise ratio. Our data are in line with this prediction (**Figure S5**). These results, therefore, highlight the importance of fine-grained MTL signals in revealing item-specific WM content.

Alongside these theoretical and empirical contributions, our data also provide additional insights into the conditions under which the MTL is relevant for WM. First, our findings suggest that the MTL’s contribution to WM does not depend on whether task demands exceed a limited WM capacity ^8^, although this account has been proposed when interpreting some recent findings for WM tasks using complex stimuli or a higher memory set size ^8,15,19–21^. Second, our analysis has focused on the mid-delay activity ^64^ and hence our findings could not be explained by the MTL’s contribution to WM retrieval ^75^. Furthermore, while our findings do not preclude the potential involvement of the MTL during perceptual encoding ^76^, perceptual involvement could not account for the results based on the comparison between the cued and uncued items ^3–5^. If the MTL primarily contributes to perceptual encoding instead of WM retention, we should have observed a comparable amount of information for both study items in the MTL, as they are presented in the same data acquisition TR before cue onset. Since participants do not know the cued item ahead of time, they need to initially remember both items. In line with this interpretation, a time-varying IEM analysis shows that aLEC-DG/CA3 indeed contains a comparable amount of information related to both the cued and uncued items at an earlier time point in the task (**Figures S6A & B**). Yet, during the mid-delay period (**Figures S6A & C**), aLEC-DG/CA3 contains significant information for the cued relative to the uncued item – in a similar way as that shown in the previous research ^3,5^. Although it is well acknowledged that the current recording method has its inferential limitations in the time domain, these data suggest that the entorhinal-DG/CA3 pathway supports the representation of a retrospectively selected memory item during a short delay – a hallmark of WM ^55,56^.

Several open questions remain to be addressed by future research. First, it is unknown how WM representation in the MTL is compared with that retained in distributed neocortical areas ^2,29,73^. Although the IEM approach allows the reconstruction of information in neural signals, it is not well-suited to directly compare information reconstruction across brain regions. Such a comparison would be complicated by several issues, including the difference in the number of voxels involved and the lack of interpretability of null results when both brain regions contain some WM information. To improve interpretability, we have used the results based on the uncued item as a within-ROI control and contrasted how information specific to the cued item (cued vs. uncued) differs between MTL ROIs and a theoretically irrelevant control region (i.e., the amygdala). One additional potential approach is to examine how the representations of remembered items are correlated across brain regions (**Figure S7A**). The rationale is that delay-period neural patterns across trials should be correlated for two brain regions containing the same information ^77^, as compared with brain regions that do not hold consistent information (**Figure S7B**). We tested this conjecture by examining the neural similarity across trials between the aLEC-DG/CA3 and a benchmark ROI in the superior temporal lobule (SPL) – a region that is consistently linked with item-specific information during visual WM retention both in the current data (**Figures S5**) and in the previous research ^4–6^. Supporting this prediction, we found that the similarity of neural patterns between the aLEC-DG/CA3 and the SPL has increased from the pre-stimulus baseline to the WM retention period (**Figure S7C**), which contrasts with the lack of changes in the correlation of across-trial neural patterns between aLEC-DG/CA3 and the amygdala control ROI (**Figure S7D**). These data suggest that the information content present in delay-period entorhinal-DG/CA3 activity shows some shared variance across trials with that in a well-recognized neocortical area related to visual WM ^4–6^. Future research with direct recordings should further investigate the fine-scale temporal dynamic underlying these similar neural patterns across brain regions during WM.

Second, it remains to be clarified how the MTL circuitry is tuned to a certain orientation, although one of the analytical tools we used was inspired by findings based on neuronal tuning properties from the visual cortex ^65,78^. This is because the assumed orientation channels in IEM do not reflect the underlying neuronal tuning properties and are interpretable only within the assumed model ^65,79^. Previous research using this method thus has primarily focused on inference related to the presence or absence of information content in the neural data ^4,5,66,67,78^, instead of properties of neural tuning. In the current study, these IEM results are supported by the less assumption-laden results based on stimulus-based representational similarity analysis ^63^. These two approaches are therefore complementary to each other.

Third, the often-neglected role of the MTL in visual processing needs to be further explored. Our findings suggest that the entorhinal-DG/CA3 pathway in the MTL may play a role in retaining of task-relevant item-specific visual WM content, which could not be attributed to perceptual processing alone. These data adds to a growing body of literature that considers the MTL as an important part of the visual system, serving functions ranging from retinotopic coding ^80^ to predictive coding ^81^. Although retinotopic coding as a form of perceptual processing could underlie WM representations for orientation information, our data highlight that the MTL is sensitive to the retrospectively selected information – a hallmark of WM ^55^. In addition to generalizing these findings from orientation to other surface features such as colors and shapes, future research should further examine how frontal-parietal mechanisms related to visual selection and attention interacts with the MTL system ^56^.

## Conclusion

In sum, our data demonstrate that the MTL’s entorhinal-DG/CA3 pathway retains precise item-specific WM information, similar to that present in other distributed neocortical areas ^4,5^. These results suggest that the same neural mechanisms underlying the fidelity of long-term episodic memory ^36,37,39–43^ is involved in representing precise item-specific WM content. Our data, therefore, provide broader insights into the fundamental constraints that govern the quality of our memory across timescales.

## Materials and Methods

### Participants

Sixteen right-handed participants (mean ± s.e.m.: 21.32 ± 0.73 years old, 8 females) were recruited for the study with monetary compensation ($20/hour). This sample size was designed to be no smaller than that involved in the prior studies using similar experimental paradigms and analytical procedures ^3–5^. All participants reported normal or corrected-to-normal visual acuity and no history of neurological/psychiatric disorders or prior psychostimulant use. They provided written informed consent before the study, following the protocol approved by the Internal Review Broad of the University of California, Riverside.

### Visual WM task

Participants performed an orientation visual working memory task adapted from previous studies ^3,5^ inside an MRI scanner (**Figure 1A**). Briefly, on each trial, we sequentially presented two sine-wave gratings (~4.5° of visual angles in radius, contrast at 80%, spatial frequency at ~1 cycle per visual degree, randomized phase) at the center of the screen. Each grating appeared for 200 ms, with a 400-ms blank screen in between. The two gratings had different orientations randomly drawn from nine predefined orientations (0 to 160° in 20° increments) with a small random angular jitter (± 1° to 5°). Following the offset of the second grating of each pair by 400 ms, we presented a cue (“1” or “2”, corresponding to the first or second grating, respectively) for 550 ms to indicate which grating orientation the participant should remember and maintain over an 8,750-ms delay period. We instructed participants to remember only the cued grating and to ignore the uncued one. After the delay period, we presented a test grating initially aligned to a random orientation. Participants then pressed the response box buttons to continuously adjust the test grating until it matched the orientation of the cued grating based on their memory. We asked the participants to make a response within 3,500 ms following the onset of the test grating (averaged median response time across participants: 2,929 ± 156 ms). After the response, we provided feedback to the participants by presenting a line marking the correct orientation, which was followed by an inter-trial interval of 3,500 or 5,250 ms. Participants completed 10 blocks of 18 trials, yielding a total of 180 trials inside the scanner. Before scanning, they completed 2 blocks of 18 trials outside the scanner for practice. The cue position and the orientations of presented gratings were randomly intermixed within each block.

Under an effective set size of one item, participants’ recall performance was high (**Figure 1B**), with most recall errors centered around ± 45° of the cued orientation (~97% of the trials) within the ± 90° range. Hence, we have retained all trials when investigating the amount of WM information in the recorded neural data during the delay period. We use the absolute recall error as a trial-level estimate of recall fidelity ^56^, assuming that large recall errors were driven by imprecise WM instead of other factors, such as occasional attentional lapses ^68^. To minimize the contamination of these factors in linking the neural data with the behavioral data, we have focused on the trials where participants have recalled within the 3 SD of the aggregated recall error distribution (**Figure 4A**; see details in a subsequent section).

### MRI Data Acquisition and Pre-processing

We acquired neuroimaging data using a 32-channel sensitivity encoding (SENSE) coil in a Siemens Prisma 3.0-Tesla scanner. We first acquired a high-resolution 3D magnetization-prepared rapid gradient echo (MP-RAGE) structural scan (0.80 mm isotropic voxels) and then functional MRI scans consisted of a T2*-weighted echo-planar imaging (EPI) sequence: TR = 1750 ms, TE = 32 ms, flip angle = 74°, 69 slices, 189 dynamics per run, 1.5 × 1.5 mm^2^ in-plane resolution with 2 mm slice thickness, FOV read = 222 mm, FOV phase = 86.5%. This sequence was optimized for high-resolution functional MRI with whole-brain coverage for the scanner. Each functional run lasted 5 minutes and 30.75 seconds. At the end of the experiment, we acquired two additional scans with opposite phases to correct for EPI distortions ^82^.

We preprocessed neuroimaging data using the *Analysis of Functional NeuroImages* (AFNI) software ^83^. Briefly, functional data were de-spiked (*3dDespiked*), slice timing corrected (*3dtshift*), reverse-blip registered (*blip*), aligned to structural scan (*align_epi_anat.py*), motion-corrected (*3dvolreg*), and masked to exclude voxels outside the brain (*3dautomask*). To avoid introducing artificial autocorrelations in later analyses, functional data were not smoothed. For the same reason, we extracted the raw BOLD signals from the middle 3 TRs of the 5-TR retention interval for later analyses without fitting the data to the hemodynamic model ^5^. These raw BOLD signals were z-scored within each block/run, before extracting the TRs of interest. In particular, we convolved the data from the 5 TR delay period with a set of weights (i.e., 0, 1, 2, 1, 0) that resembled the TENT function in AFNI to maximize the inclusion of mid-delay activity for later analysis ^64^. This approach factors in 5-6 s of hemodynamic adjustment ^84^ and has been considered fundamentally conservative in estimating delay-period activity ^85^. This approach also provides a reasonable estimate for the BOLD response around a given TR with an improved signal-to-noise ratio without assuming the shape of the underlying hemodynamic response ^86^. We also performed the time-varying version of this analysis by shifting the peak of the TENT function over time (see **Figure S6** for details).

To retain the consistency with the prior research, we defined participant-specific MTL ROIs (bilateral hippocampal DG/CA3, CA1, and subiculum, entorhinal/perirhinal cortex, and parahippocampus, see **Figure 2A**) based on the T1 image using the same segmentation algorithm from the previous studies ^53,60^. In brief, using the *Advanced Normalization Tools* ^87^, this algorithm aligns an in-house segmented template to each participant’s T1 image. This template contains manually labeled ROIs for hippocampal subfields (DG, CA3, CA1, subiculum) and other verified MTL subregions (aLEC, pMEC, perirhinal, and parahippocampus). The efforts to select and verify these MTL ROIs have been detailed in previous studies ^53,88^. In brief, in addition to the commonly identified perirhinal and parahippocampus ROIs, hippocampal subfields were manually identified and aggregated from a set of T1 and T2 atlas images based on prior harmonized efforts ^89^. Entorhinal ROIs (aLEC and pMEC) were added to the template from a previous study ^90^. For functional analysis, we combined DG and CA3 subfields as a single label given the uncertainty in separating signals from them in fMRI data ^60^. In addition, we also verified our findings in hippocampal subfields based on a different segmentation protocol via *FreeSurfer* ^91^, which yielded consistent findings (**Figure S8**). Therefore, our current observations are unlikely to be limited to a specific parcellation procedure of hippocampal subfields.

Furthermore, we identified subject-specific segmented amygdala as a control ROI based on participant-specific *Freesurfer* parcellation ^92^. The amygdala is a part of the limbic system traditionally considered a central brain region processing emotion-laden information. Because the task stimuli (orientation gratings) and testing procedure (no reward manipulation) in the current study are emotionally neutral, the amygdala is therefore theoretically irrelevant for the current study ^61^. Furthermore, as its signal-to-noise ratio is similar to adjacent structures, the amygdala can serve as a control site for the observation in other MTL ROIs.

### Stimulus-based Representational Similarity Analysis

To examine whether MTL delay-period activity can distinguish different cued orientation gratings, we performed a stimulus-based representational similarity analysis ^93^. The rationale is that if the recorded neural data contain information to allow fine discrimination of the cue item, the neural data should track the feature distance between any pair of cued items across trials to allow fine discrimination of these items ^63^. Hence, we first calculated the stimulus similarity pattern across trials using 180 minus the absolute angular distance between the orientation labels of every two trials (**Figure 2B**, top panel). Next, we calculated the cosine similarity of the delay-period neural signals ***B*** across *n* voxels from the middle 3 TRs in every pair of trials (***Fi*gure 2B**, bottom panel). This yields a trial-by-trial matrix in which the similarity between voxel response vectors *B_i_* and *B_j_* from the lower diagonal. Their similarity is calculated as,

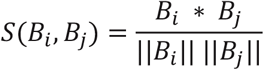

Finally, we correlated (rank-order) the neural similarity pattern and stimulus similarity pattern across trials to gauge how the recorded neural signals track the stimulus features across trials.

### Inverted Encoding Modeling (IEM)

To decode item-level information from the raw BOLD signals ^5^, we first constructed a linear encoding model to represent orientation-selective responses in multi-voxels of activity from a given brain region. We did not impose any additional feature selection procedures other than using the anatomically defined ROIs to identify relevant multi-voxel features in this analysis (see **Table S3** for the number of voxel/features included for each subject in each ROI). We assumed that the response of each voxel is a linear summation of 9 idealized information channels (**Figure 2B**), estimated by a set of half-wave rectified sinusoids centered at different orientations based on the tuning profile of orientation-sensitive neural populations. Hence, we formalized the observed raw BOLD signals ***B*** (*m* voxels × *n* trials) as a weighted summation of channel responses ***C*** (*k* channels × *n* trials), based on the weight matrix, ***W*** (*m* voxels × *k* channels), plus residual noise (***N***),

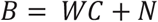

Given ***B_1_*** and ***C_1_*** from a set of training data, the weight matrix can be calculated as,

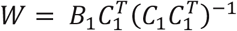

The training weight matrix ***W*** is used to calculate a set of optimal orientation filters ***V***, to capture the underlying channel responses while accounting for correlated variability between voxels (i.e., the noise covariance), as follows,

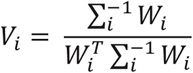

where 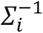 is the regularized noise covariance matrix for channel *i* (1 to 9), estimated as,

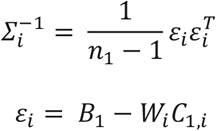

Here, *n_1_* is the number of training trials, and *ε_i_* is a matrix residual based on the training set ***B_1_*** and is obtained by regularization-based shrinkage using an analytically determined shrinkage parameter. Next, for the independent hold-out test dataset ***B_2_***, trial-by-trial channel responses ***C_2_*** are calculated as follows,

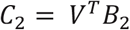

We used a leave-one-out cross-validation routine to obtain reliable estimate channel responses for all trials. For each participant, in every iteration, we treated all but one block as ***B_1_*** and the remaining block as ***B_2_*** for the estimation of ***C_2_***. This analysis yielded estimated channel responses ***C_2_*** for each trial, which were interpolated to 180° and circularly shifted to a common center (0°, by convention). We reconstructed these normalized channel responses separately using orientation labels of the cued item, the uncued item, and shuffled orientations. We then quantified the amount of item-related information (*R*) by converting the average channel response (*z*) to polar form given *ψ* as the vector of angles at which the channels peak (*z* = *Ce*^2*iψ*^). We then projected them onto a vector with an angle of 0°,

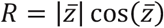

With whole-brain coverage, we performed an additional searchlight procedure in combination with the IEM analysis to replicate the previous findings ^5^. First, we normalized participants’ brain data to an MNI template using the *Advanced Normalization Tools*. Second, we defined a spherical “neighborhood” (radius 8.0 mm) centered on voxels in a cortical mask containing only gray matter voxels. We discarded neighborhoods with fewer than 100 voxels. Last, we estimated item-related information (*R*) about the to-be-remember item based on the IEM analysis outlined above to assess WM information within each searchlight sphere. We obtained consistent findings as compared with the previous findings (**Figure S3**), suggesting the reliability of the current data.

### Linking IEM Reconstruction with Behavioral Recall Performance

To examine how the IEM reconstruction of the cued item in the aLEC-DG/CA3 pathway is associated with recall fidelity, we performed the IEM analysis based on data combined from the aLEC and DG/CA3 ROIs. Similar to the analytical framework outlined above, we split each participant’s data into random blocks of 18 trials and then perform a leave-one-block-out analysis to obtain IEM reconstructions for all trials in each block based on the weights trained from other blocks. As this analysis is agnostic to participants’ recall performance at this stage, if IEM reconstruction is not associated with participants’ recall fidelity, the reconstructed information channels should be comparable regardless of recall errors. To test against this prediction, we split participants’ data into small- and larger-error trials. First, as the angular resolution is at least 20° for any two items in the current design, we defined small-recall error trials as those in which participants have reported within one similar item away (absolute recall error < 20°; 149 ± 3 trials). Next, to separate larger-recall errors based on less precise WM representation from those attributable to attention lapses ^68^, we adopted a widely-used thresholding heuristic to find potentially different categories of data points based on the empirical SD of a distribution. Specifically, in our current data, we first calculated the empirical SD (17.33°) of the aggregated raw recall error distribution from all subjects across 2880 trials (ranging from −90° to +90°), which captures the overall variability in participants’ recall performance without *a priori* model assumption. We then retained the larger-recall error trials within 20° to 3 SD of the recall error distribution (27 ± 3 trials; **Figure 4A**). These larger-error trials presumably contain mostly imprecise recall responses, instead of infrequent extra-large errors that could be attributed to other factors like attentional lapses ^68^. Considering that most of the trials have a recall error of ± 45° out of the ± 90° range in every subject by visual inspection (97% of the trials, **Figure 1B**), we have also used 45° of absolute recall error as a cut-off for extra-large error trials and obtained similar findings in subsequent analyses (see **Supplementary Information**). To ground our analysis in empirical data, we therefore have focused on the results based on the SD thresholding heuristic in the current report.

To balance the trial counts between these two categories of trials, we resampled the same number of trials based on the number of larger-error trials from the small-error trials for 5,000 times. This resampling procedure ensures that the average IEM reconstruction from the small-error trials is estimated based on the same number of trials as compared with the larger-error trials – an approach often used to obtain less biased estimates of neural measures across different behavioral trial types ^24,69,70^. We contrasted the difference in IEM reconstructions for the cued item in the aLEC-DG/CA3 between these two categories of trials across participants.

### Statistical Procedures

We evaluated statistical significance based on conventional within-subject statistical procedures, such as paired-sample t-tests, with two-tailed p values. We estimated the size of these effects based on Cohen’s *d*. Except for pre-defined contrast analysis (e.g., cued vs. uncued), we corrected for multiple comparisons by using Bonferroni correction with an alpha level set as 0.05 ^94^. For visualization of variability in mean estimates, we have used the standard error of the mean across participants (s.e.m.), namely the standard deviation of a measure divided by the square root of sample size, as error bars (or areas) in Figures 2, 3, and 4.

## Data and Code Availability

Non-identified data used in this study regarding MTL activities across ROIs and trial-by-trial behavior responses are available via the Open Science Framework repository (https://osf.io/zvdnr/). Custom code that supports the findings of this study is available from W.X. upon request.

## Acknowledgments

We thank all the participants who selflessly volunteered their time for this study. We thank Nicholas J. Tustison and Jason Langley for their technical support. This work was supported by the National Institute of Mental Health (1R01MH117132, PI: W. Z.). W.X. was funded by the National Institute of Neurological Disorders and Stroke Competitive Postdoctoral Fellowship Award.

## Author contributions

W. X. and W. Z. conceptualized the study and wrote the paper. W. X., M. C. contributed to data collection. E. E., M. Y., and K. Z contributed to data analysis, interpretation, and/or manuscript preparation.

## Supplementary Information

### Association between recall fidelity and IEM reconstruction

In addition to using an empirical criterion to separate in-memory trials from those extra-large error trials susceptible to occasional attentional lapses ^68^, we have also tried another thresholding heuristic. As shown in **Figure 1A**, most trials from each participant fall within this 45° of absolute recall error (i.e., half of the 90° range), and the trials larger than this number are rare (~5 out of 180 trials). We, therefore, used 45° of absolute recall error as a cut-off to identify the imprecise recall trials that are greater than 20° but smaller than 45° of absolute recall error.

We performed the same analysis to obtain trial-by-trial IEM reconstructions based on delay-period BOLD signals aggregated from the aLEC and DG/CA3, and then resampled the same number of trials to estimate the IEM reconstructions for the small-error and larger-error trials (<20° vs. 20° - 45° of absolute recall error). Consistent with the 3-SD heuristic, we found that the small-error trials yielded significant IEM reconstructions for the cued item (t(15) = 4.34, p = 5.74e-04, Cohen’s d = 1.12), whereas the larger-error trials did not (t(15) = −0.69, p = 0.50, Cohen’s d = −0.18). We then contrasted the difference in IEM reconstructions between these small- and large-error trials across participants. We found that IEM reconstruction for the cued item from the combined aLEC-DG/CA3 has better quality in the small-error trials, as compared with that in the larger-error trials (t(15) = 3.41, p = 0.004, Cohen’s d = 0.88). These results suggest that higher-quality visual WM content in the entorhinal-DG/CA3 pathway during the delay period is associated with better subsequent recall fidelity and that this association is robust to the selection of the cut-off for extra-large recall errors.

**Figure S1.**
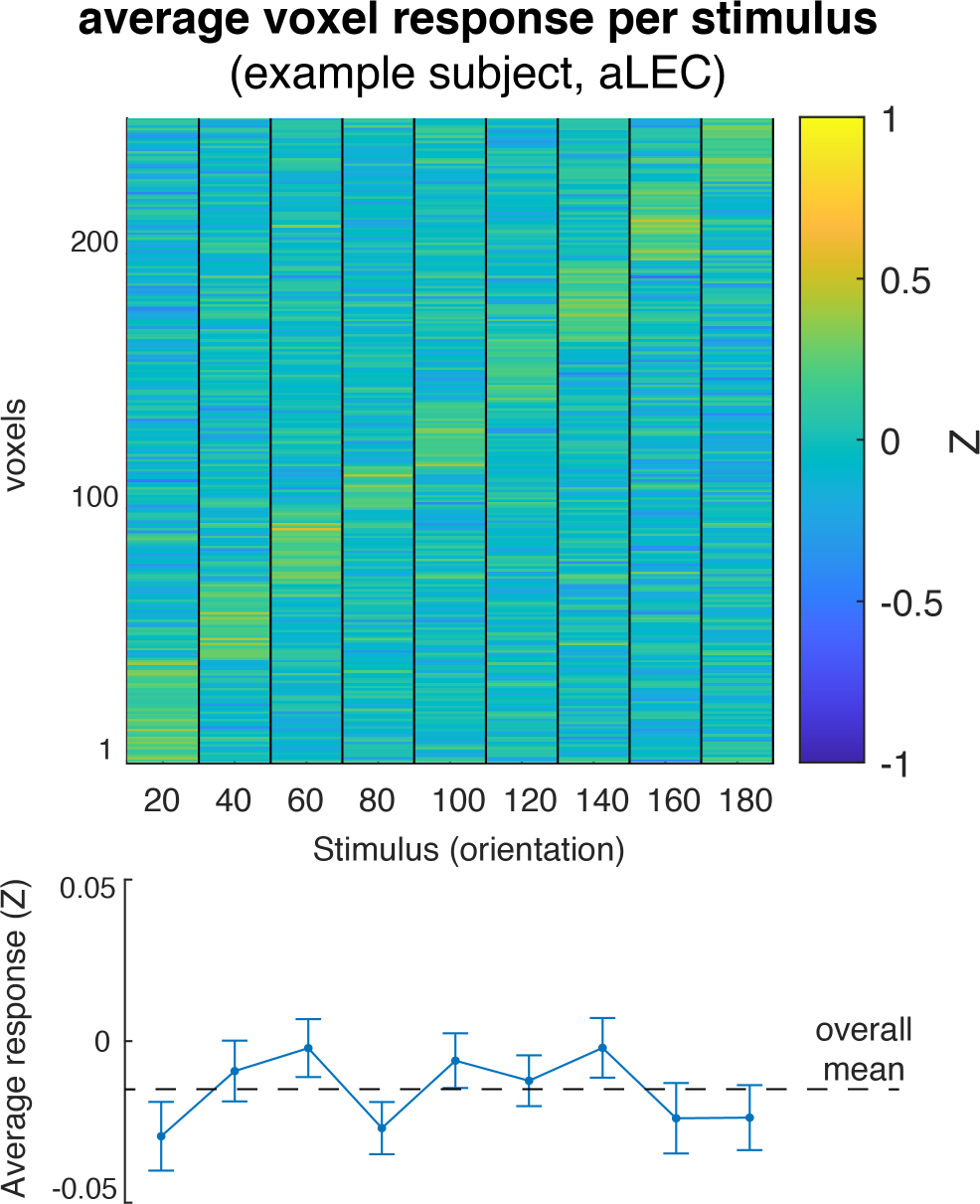
Voxel responses in an example ROI (aLEC) for different remembered stimuli from one example subject. We sorted these voxels based on the magnitude of BOLD response to different orientation stimuli. This analysis only serves illustrative purposes. The reliability of these multi-voxel patterns can be examined based on stimulus-based representational similarity analysis as detailed in the main text.

**Figure S2.**
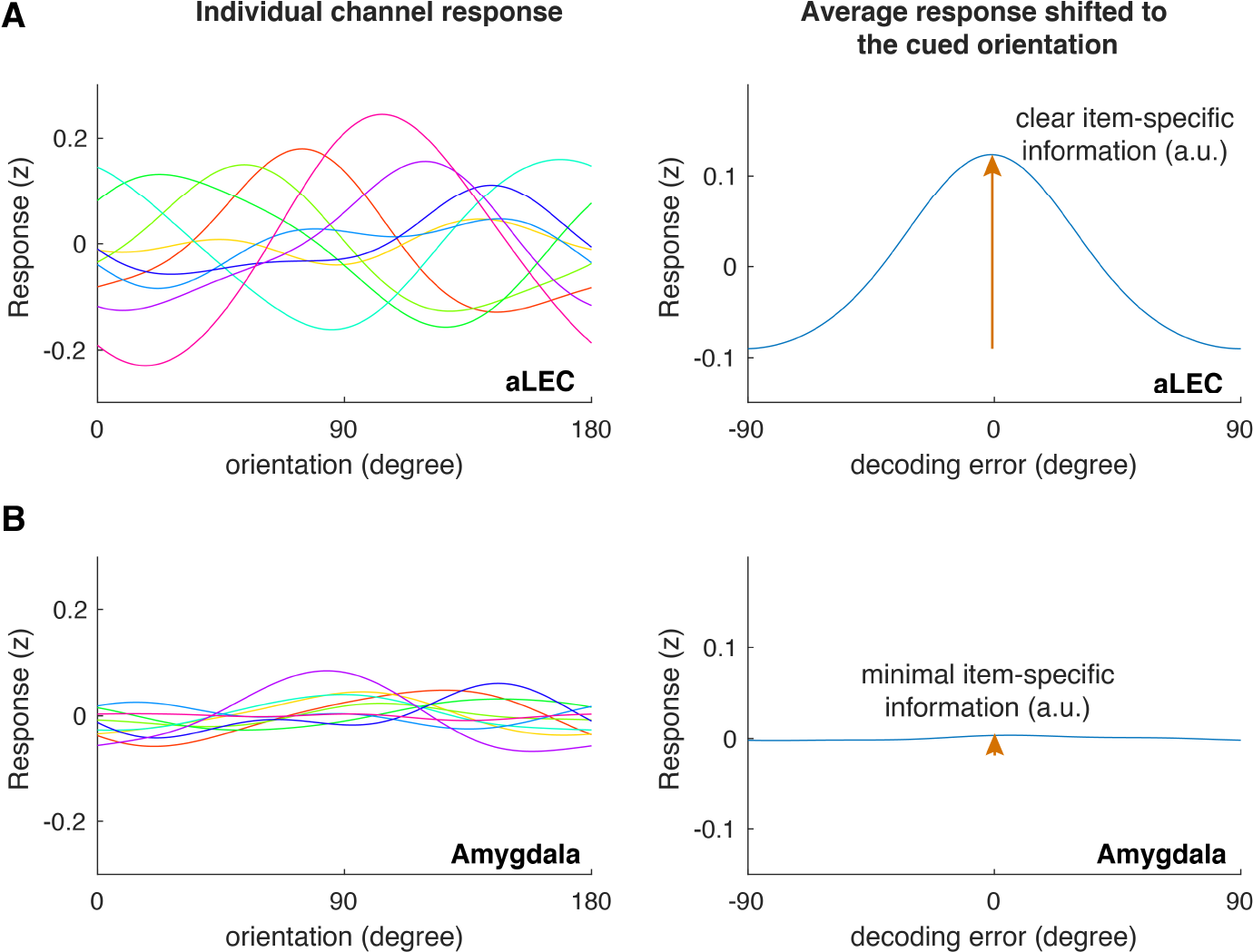
Example channel responses before and after shifting to the cued orientation for aLEC (A) and the amygdala (B). Based on a level-one-block-out cross-validation approach, we reconstructed the assumed neural channel response model reported in **Figure 3B**. Before shifting individual channel responses to the cued orientation, it is expected that these tuning responses should show separate peaks across the feature space (right panels). After shifting individual channel responses to the cued orientation, if there is information about the cued orientation assumed by the model, it is expected that the average channel response should peak and center around a 0-degree error (left panels). In cases where information is unrelated to the encoded orientation (e.g., amygdala), the IEM approach is expected to fail to capture and reconstruct meaningful orientation information because the unrelated noise can distort the training weights of the encoding model. Note that the results after shifting the channel responses to the cued orientation are the same as that shown in **Figure 3**.

**Figure S3.**
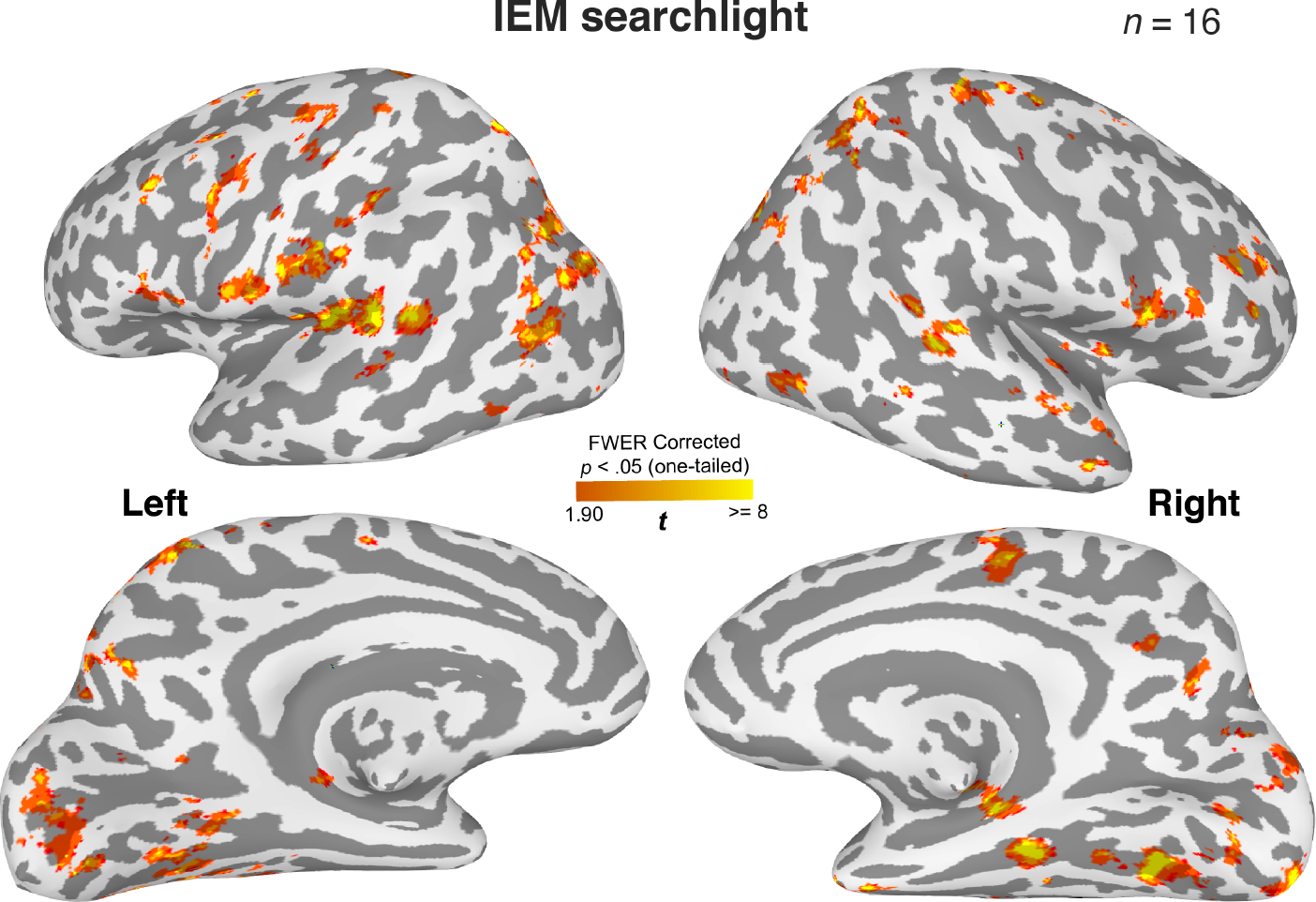
Distributed brain regions retain information about the cued item during WM. A roving searchlight procedure was combined with the inverted encoding modeling (IEM) to identify brain regions containing item-specific WM content for the cued item ^5^. This analysis shows that distributed brain regions retain decodable item-specific information for the cued orientation, replicating the previous findings ^3–5,29^. Cluster-based correction for statistical significance: *p* < .05 (one-tail) with > 400 voxels estimated based on *3dClustSim* from AFNI. In this analysis, we have also observed significant clusters of voxels in the MTL. However, this observation is limited to small cortical surface areas. One possibility is that the 8-mm searchlight sphere may have failed to take into account the complex folding structures in the MTL, such that heterogeneous information is included for the decoding analysis. Consequently, the contribution of item-specific WM information in certain MTL voxels can be attenuated. We tested this prediction by aggregating the data from the whole hippocampus across subfields and summarized the findings in **Figure S5**, which supports our prediction.

**Figure S4.**
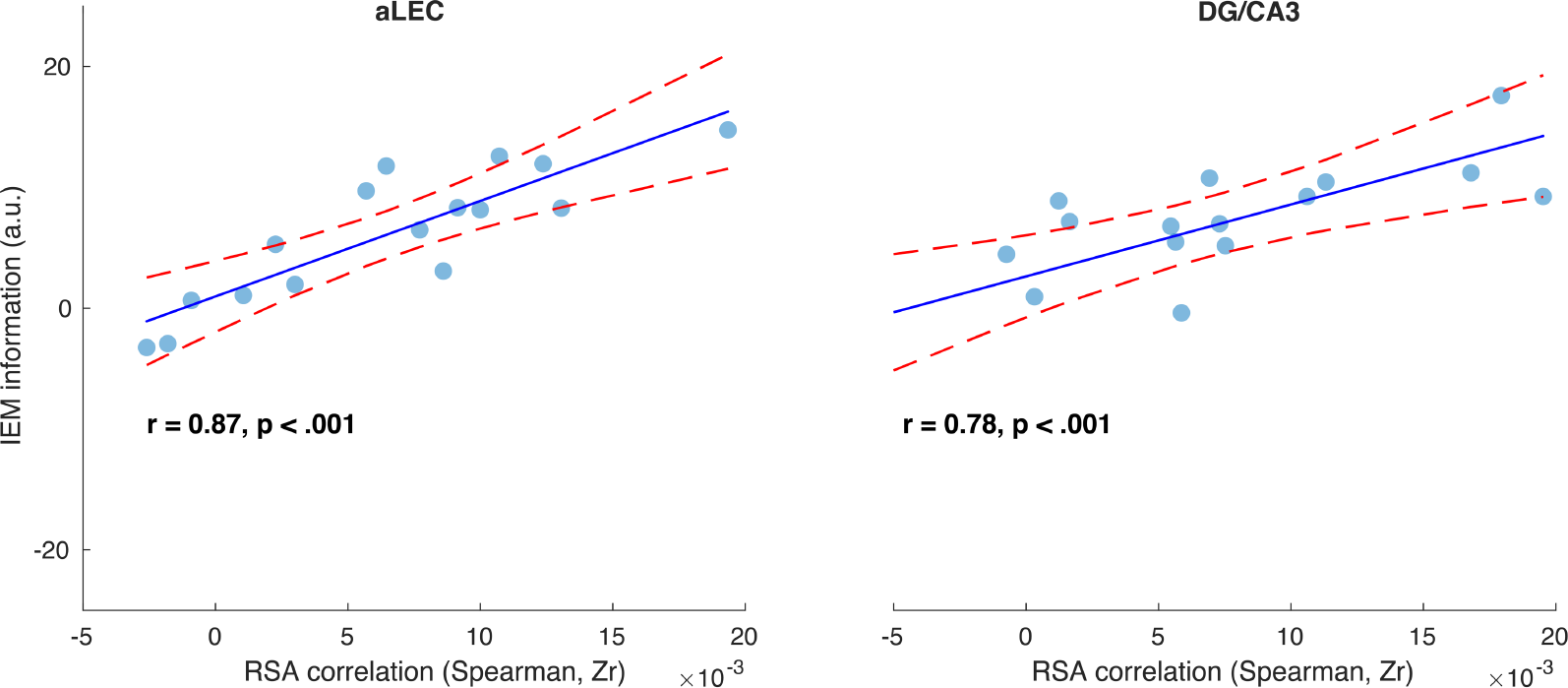
Stimulus-based representational similarity analysis (RSA) and inverted encoding model (IEM) reveal shared item-related variance in the observed neural data. In both the aLEC and DG/CA3 ROIs, the association between the patterns of the task stimuli and neural responses (RSA) was highly correlated with IEM decoding performance across participants (aLEC: *r* = 0.87, *p* < .001; DG/CA3: *r* = 0.78, *p* < .001), even though these two methods have different assumptions and analytical procedures. This observation suggests that item-specific WM content in MTL regions can be reliably captured by different analytical procedures. The x-axis shows the values of the RSA correlation between trial-by-trial stimulus similarity patterns and observed neural similarity patterns of fMRI BOLD activity. Data on the y-axis reflect the resultant vector length of the normalized reconstructed orientation information channels based on IEM analysis from the same region. Individual points represent the results from individual participants. The solid lines are linear fits of the data, and the dashed lines are 95% confidence intervals of the linear fits.

**Figure S5.**
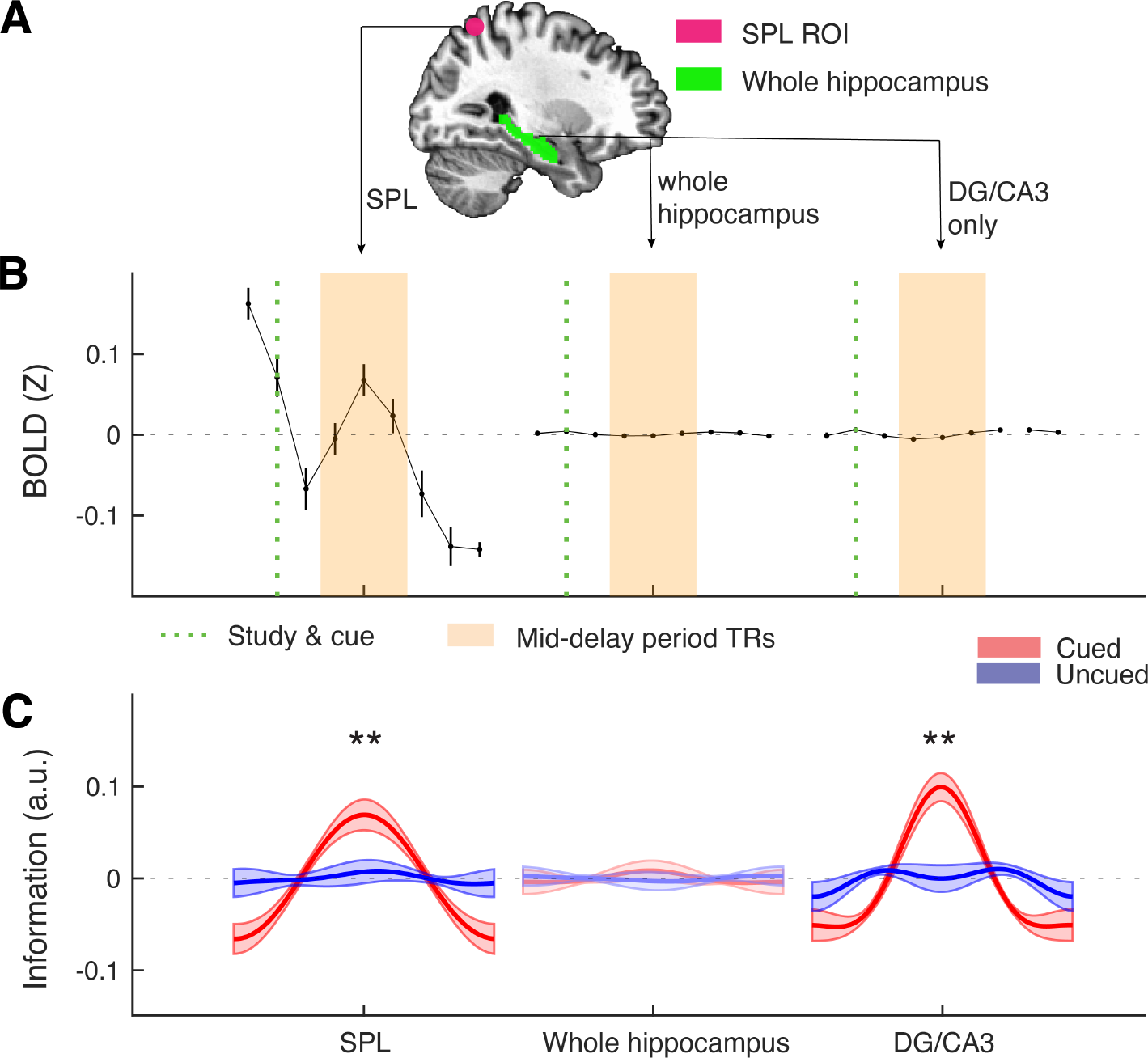
Analyses based on the whole hippocampus, as compared with a benchmark sphere ROI in the posterior parietal cortex (e.g., superior parietal lobule, SPL) and the hippocampal DG/CA3 subfield. (**A**) As the posterior parietal cortex has consistently implicated to support visual WM representations ^4,5^, we identified the local peaks of the bilateral posterior parietal clusters based on the searchlight analysis in **Figure S3** (MNI coordinate: left, x = −16, y = −64, z = 58; Right, x = 30, y = −56, z = 58) to extract two 8-mm sphere ROIs from the regions. These ROIs fall within the SPL, with their central coordinates with those from a previous study ^5^ based on similar methods (e.g., left, x = −19, y = −63, z = 55; right, x = 20, y = − 58, z = 57). For visualization, we plotted one of the spheres in the figure, in combination with the hippocampus (right brain). (**B**) Raw BOLD signals in each voxel were z-scored over time separated in each block before extracting the trial structures and then averaged for each ROI. Mid-delay 3 TRs are represented by the shaded area in orange. Consistent with the previous observations ^4,5,7^, the SPL in the posterior parietal cortex shows robust BOLD modulation during the visual WM task. In contrast, the hippocampus as a whole and its relevant subfield – DG/CA3, do not show the same magnitude of BOLD modulation as the SPL. (**C**) While both the SPL and DG/CA3 retain precise item-specific information about the cued item relative to the uncued item, the whole hippocampus however does not show this pattern, suggesting that the inclusion of heterogeneous voxels from CA1 and subiculum may affect IEM performance. This is unsurprising because the current IEM analysis did not include additional feature selection procedures, and hence the inclusion of uninformative voxels would make the weights trained from these data less information, compromising the subsequent IEM reconstruction. Error bars or shaded areas represent the standard error of the mean (s.e.m.) across participants. **p. < .01; a.u. = arbitrary unit.

**Figure S6.**
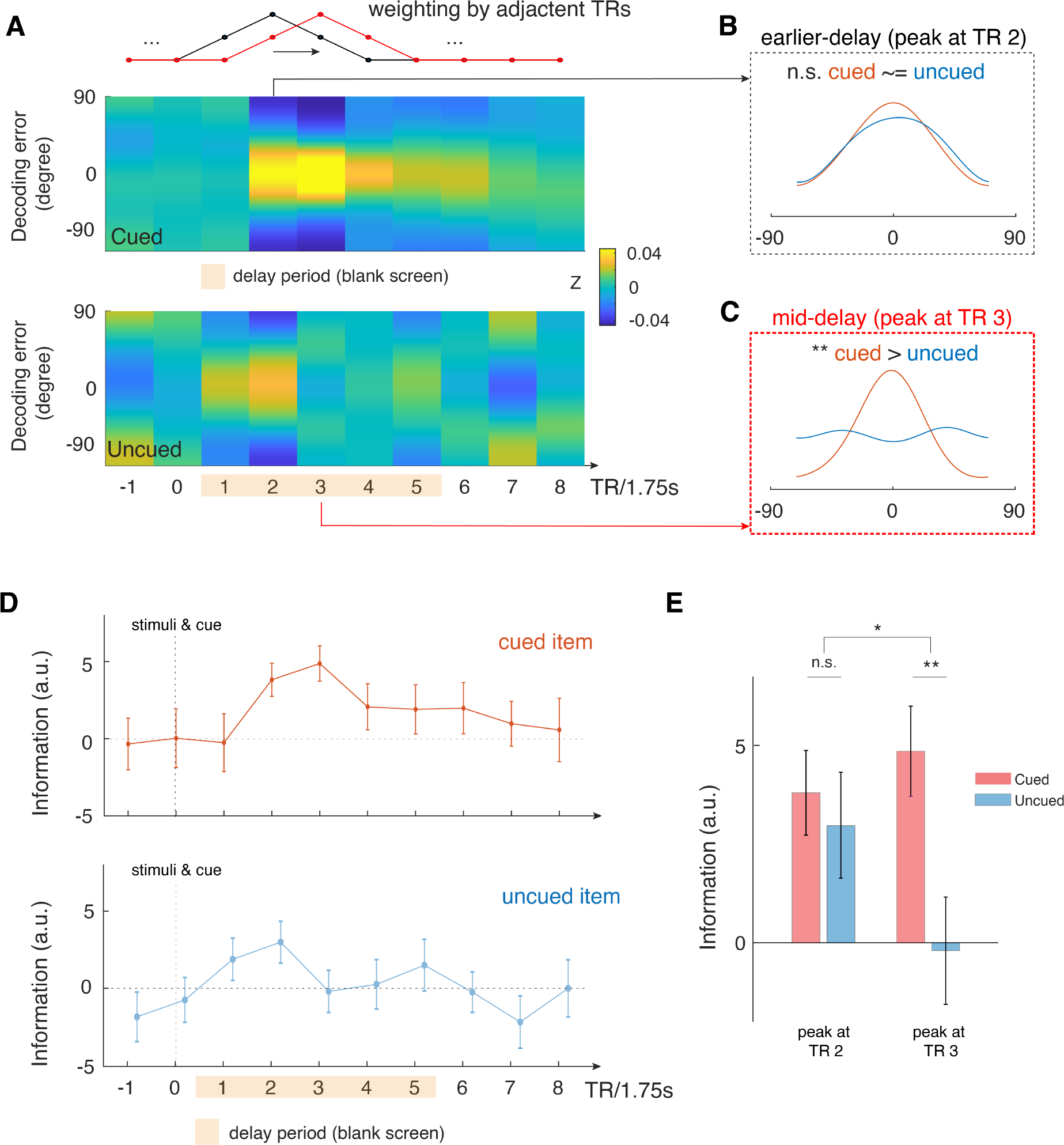
Time-varying IEM analysis shows that mid-delay period activity in aLEC-DG/CA3 contains item-specific information that could not be attributed to perceptual processing alone. (**A**) We performed a time-varying IEM analysis for the combined aLEC-DG/CA3 ROI based on raw BOLD signals weighted by adjacent TRs, with a moving window in steps of 1 TR. This analysis was done separately using trial labels of the cued and uncued items. TR 0 contains the presentation of study items and the retro-cue. (**B**) We find that earlier TR in the delay period (e.g., TR 2) contains information for both the cued and uncued items. (**C**) Yet, mid-delay period activity (e.g., TR 3, ~5.25s after stimulus offset) contains the most information related to the cued item, but not to the uncued items, suggesting that perceptual processing could not account for these results. (**D**) Timing-varying analysis shows that information related to the cued item increases and peaks at the mid-delay period, with attenuated information throughout the rest of the delay period. In contrast, information related to the uncued item increases after stimulus offset but dissipates afterwards. (**E**) We observed a significant interaction effect in the reconstructed IEM information between time period (earlier, TR 2 vs. mid-delay, TR 3) and cue condition (cued vs. uncued; F(1, 15) = 4.85, p < .05). These results suggest that there is additional information in the mid-delay activity related to the retrospectively selected item, which could not be accounted for perceptual processing of the presented stimuli. Error bars represent the standard error of the mean (s.e.m.) across participants. \**p* < .05; \*\**p* < .01; n.s. = not significant.

**Figure S7.**
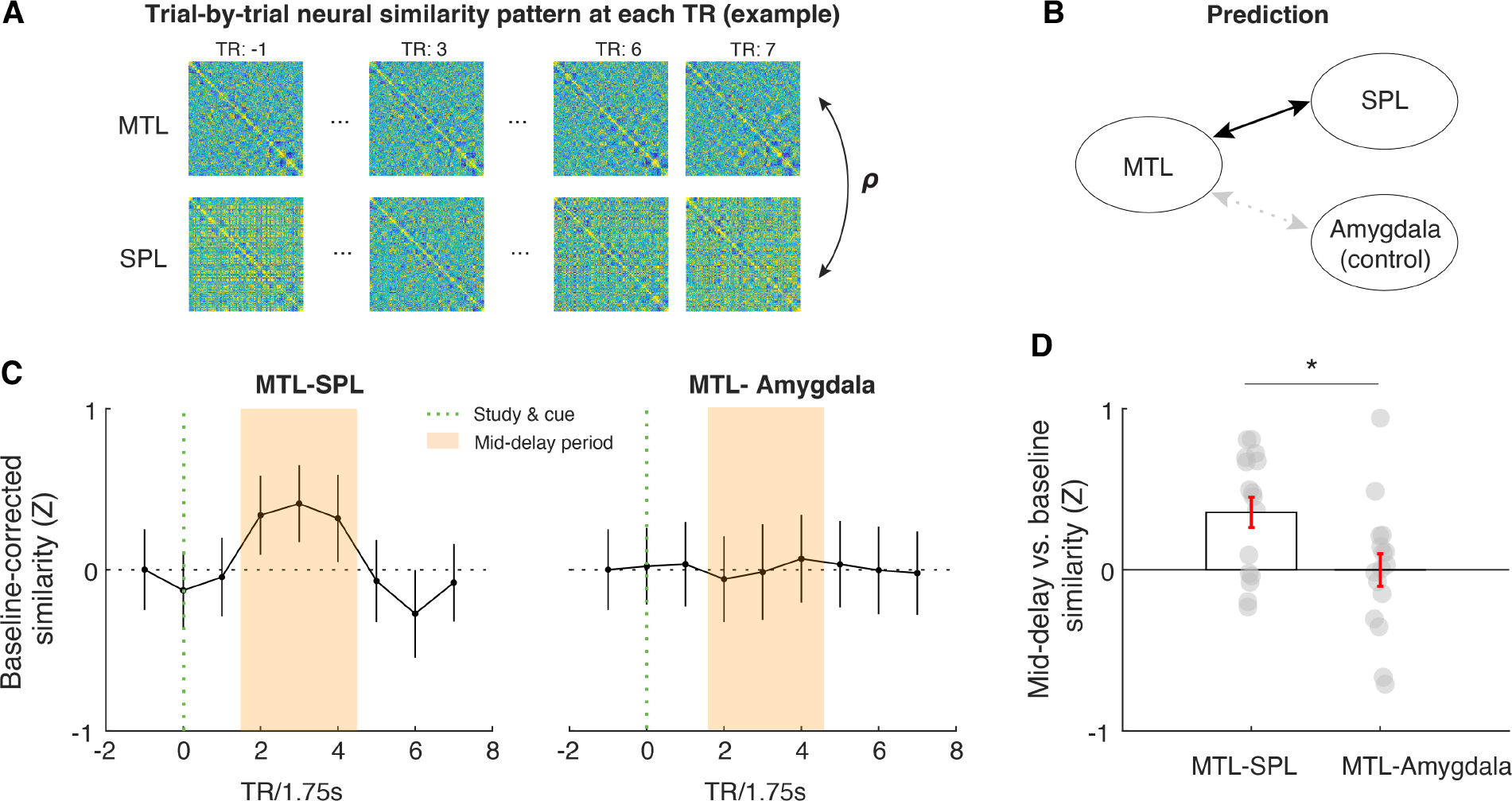
Across-region neural similarity analysis using the combined aLEC-DG/CA3 as an MTL seed region, the SPL ROI as a benchmark region, and the amygdala as a control region. (**A**) Trial-by-trial neural similarity pattern at each TR can be calculated to reflect the representational pattern within a given ROI. The similarity of these neural representational patterns, therefore, can inform us whether there is shared variance in the information content represented in different ROIs ^77^. (**B**) Because the MTL and SPL both retain information about the cued item, it is expected that their activity patterns evoked by the cued item should be similar to each other, as compared with the similarity between MTL and the amygdala control ROI. (**C**) We performed the proposed analysis at each TR. By normalizing the observed neural similarity values using the mean and standard deviation of neural similarity measures at −1 TR across participants. Following this normalization procedure, changes in the neural similarity between ROIs could not be accounted for by intrinsic neural similarity at baseline. This comparison allows us to gauge the extent to which a set of brain regions retains similar information content that is different from the similarity in overall neural signals triggered by the presentation of the same task stimuli ^77^. We find that the similarity between MTL and SPL activity patterns increases from baseline to WM retention. (**D**) Critically, this increase is absent in the similarity between MTL and amygdala control ROIs, which was supported by a significant time period (average value in middle 3TRs vs. baseline TR) and ROI (MTL-SPL vs. MTL-Amygdala) interaction effect in neural similarity measures (F(1,15) = 6.38, p < .05). These results, therefore, suggest that the DG/CA3-aLEC circuitry in the MTL shares similar information content as that in SPL during WM, which is not simply driven by the similarity of neural signals across regions induced by task stimuli. Error bars represent the standard error of the mean (s.e.m.) across participants. Grey dots in (**D**) represent data from individual subjects. *p. < .05.

**Figure S8.**
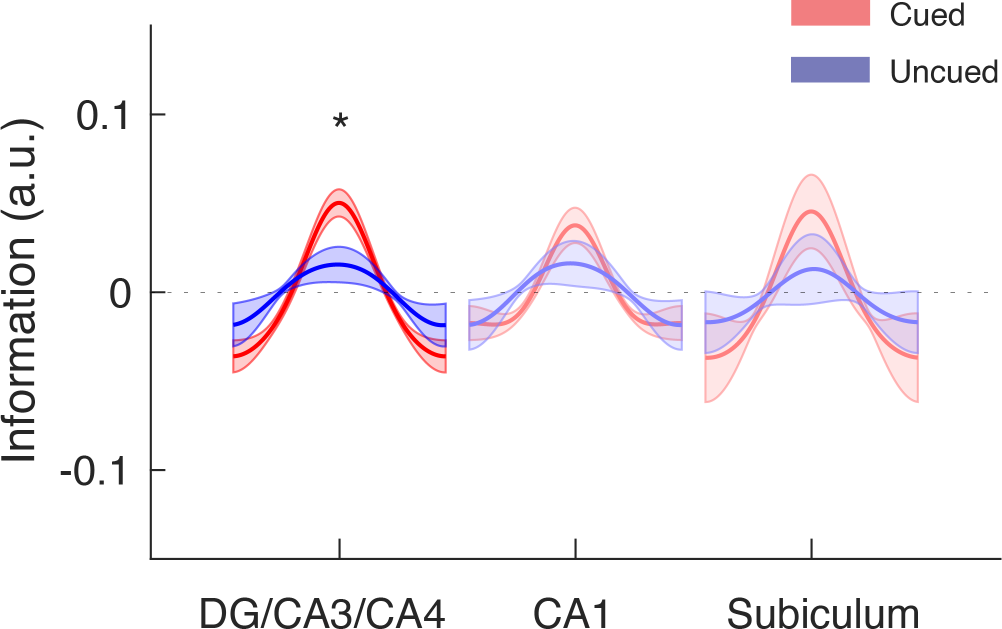
Modeling results of the hippocampal subfields based on FreeSurfer labels. In addition to the in-house subfield parcellation, we verified our findings based on the hippocampal subfield labels extracted from each participant’s MRI scan using *FreeSurfer 6.0* (https://surfer.nmr.mgh.harvard.edu/fswiki/HippocampalSubfieldsAndNucleiOfAmygdala). In this program, the DG subfield is aborted by the CA4 label. Despite different quantification methods, our findings of greater item-specific information related to the cued item in the mid-delay TRs remain statistically significant in the DG/CA3 subfield (p < .001), which is significantly greater than that for the uncued item (p < .05). In contrast, no significant difference was found for the hippocampal CA1 or subiculum subfield (p’s >.010). Shaded areas represent the standard error of the mean (s.e.m.) across participants. These results suggest that our observation in the hippocampal DG/CA3 subfield is not limited to a particular subfield parcellation method. *p. < .05.

**Table S1.** Tests of Statistical Significance in the Neural Similarity across Trials Captured by the Similarity of the Cued item *Note*: n.s. = non-significant. It is well-acknowledged that multiple comparisons would inflate TYPE-I error, and hence we adopted a relatively conservative correction procedure (Bonferroni correction^94^). This procedure reveals the significance test outcomes that could not be attributed to chance alone. Yet, it does not imply that the non-significant results indicate no effect. Rather, as shown in the table, non-significant effects are often associated with an attenuated effect size in the same direction, indicating unstable estimate across participants. We therefore focus on the regions where robust effects have been identified across participants after the correction of multiple comparisons (i.e., aLEC and DG/CA3).

|  | t | df | Cohen's d | p-uncorrected | p-bonferroni |
| --- | --- | --- | --- | --- | --- |
| aLEC | 4.29 | 15 | 1.11 | 0.0006 | **0.0052** |
| pMEC | 2.16 | 15 | 0.56 | 0.047 | n.s. |
| Perirhinal | 3.00 | 15 | 0.78 | 0.0089 | n.s. |
| Para-hippocampus | 2.70 | 15 | 0.70 | 0.017 | n.s. |
| DG/CA3 | 3.65 | 15 | 0.94 | 0.0024 | **0.019** |
| CA1 | 1.56 | 15 | 0.40 | 0.14 | n.s. |
| Subiculum | 2.53 | 15 | 0.65 | 0.023 | n.s. |
| Amygdala | 0.06 | 15 | 0.02 | 0.95 | n.s. |

**Table S2.**
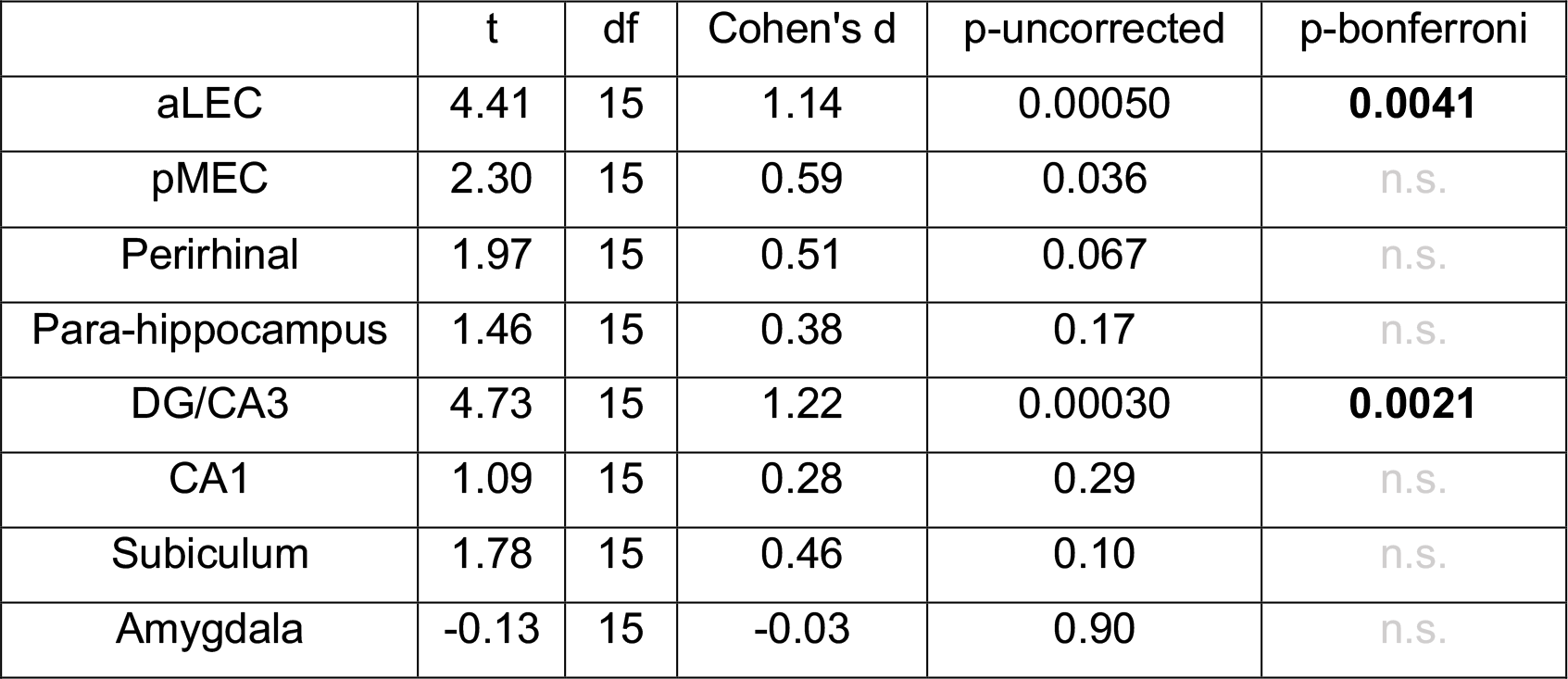
Tests of Statistical Significance in the IEM results for the Cued item *Note*: n.s. = non-significant. It is well-acknowledged that multiple comparisons would inflate TYPE-I error, and hence we adopted a relatively conservative correction procedure (Bonferroni correction^94^). This procedure reveals the significance of test outcomes that could not be attributed to chance alone. Yet, it does not imply that the non-significant results indicate no effect. Rather, as shown in the table, non-significant effects are often associated with an attenuated effect size in the same direction, indicating an unstable estimate across participants. We therefore focus on the regions where robust effects have been identified across participants after the correction of multiple comparisons (i.e., aLEC and DG/CA3).

**Table S3.**
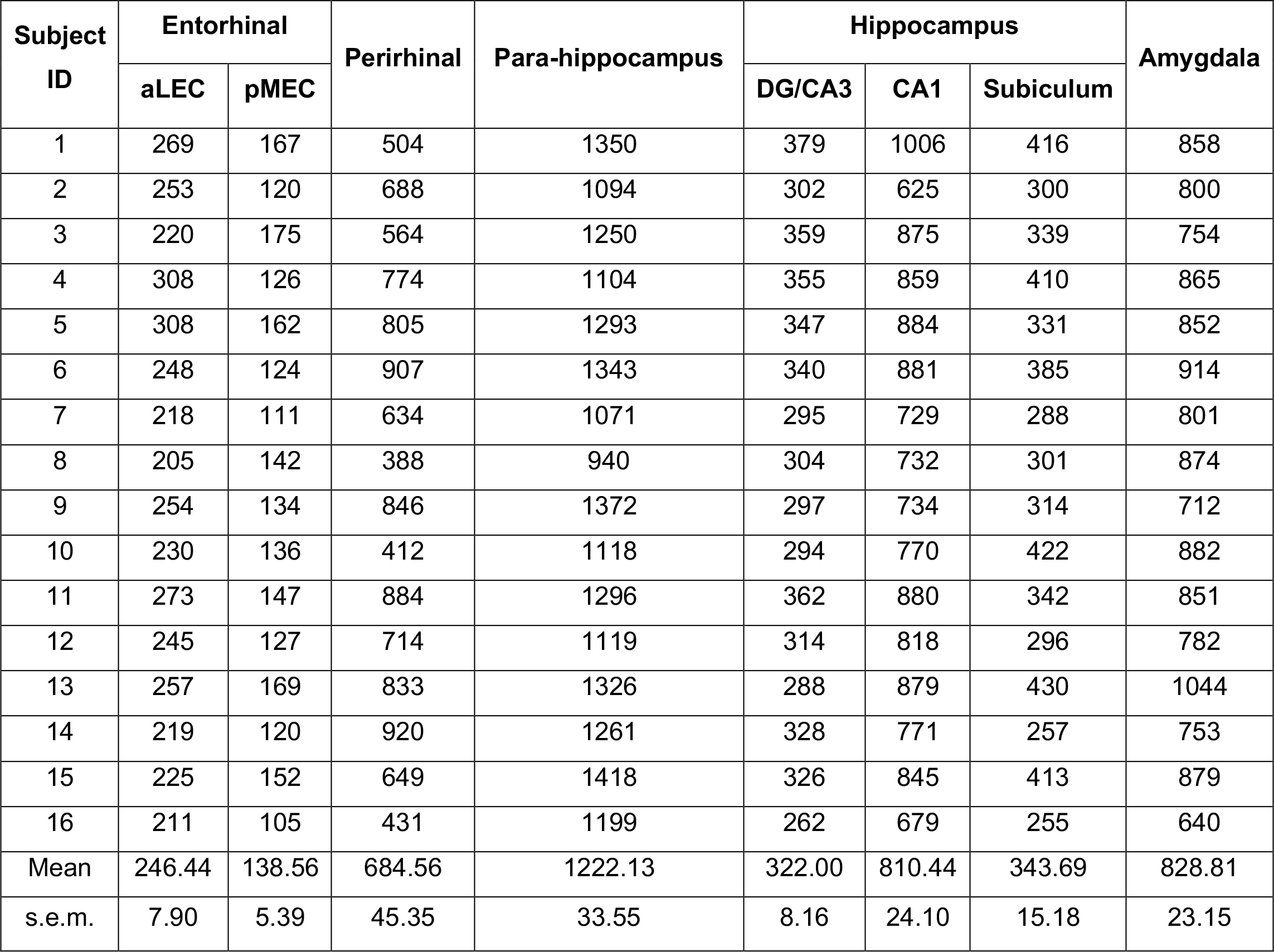
The number of voxels included for each ROI in each subject. Note: s.e.m. = standard error, which is the standard deviation divided by the square root of sample size.

| Subject ID | Entorhinal |  | Perirhinal | Para-hippocampus | Hippocampus |  |  | Amygdala |
| --- | --- | --- | --- | --- | --- | --- | --- | --- |
|  | aLEC | pMEC |  |  | DG/CA3 | CA1 | Subiculum |  |
| 1 | 269 | 167 | 504 | 1350 | 379 | 1006 | 416 | 858 |
| 2 | 253 | 120 | 688 | 1094 | 302 | 625 | 300 | 800 |
| 3 | 220 | 175 | 564 | 1250 | 359 | 875 | 339 | 754 |
| 4 | 308 | 126 | 774 | 1104 | 355 | 859 | 410 | 865 |
| 5 | 308 | 162 | 805 | 1293 | 347 | 884 | 331 | 852 |
| 6 | 248 | 124 | 907 | 1343 | 340 | 881 | 385 | 914 |
| 7 | 218 | 111 | 634 | 1071 | 295 | 729 | 288 | 801 |
| 8 | 205 | 142 | 388 | 940 | 304 | 732 | 301 | 874 |
| 9 | 254 | 134 | 846 | 1372 | 297 | 734 | 314 | 712 |
| 10 | 230 | 136 | 412 | 1118 | 294 | 770 | 422 | 882 |
| 11 | 273 | 147 | 884 | 1296 | 362 | 880 | 342 | 851 |
| 12 | 245 | 127 | 714 | 1119 | 314 | 818 | 296 | 782 |
| 13 | 257 | 169 | 833 | 1326 | 288 | 879 | 430 | 1044 |
| 14 | 219 | 120 | 920 | 1261 | 328 | 771 | 257 | 753 |
| 15 | 225 | 152 | 649 | 1418 | 326 | 845 | 413 | 879 |
| 16 | 211 | 105 | 431 | 1199 | 262 | 679 | 255 | 640 |
| Mean | 246.44 | 138.56 | 684.56 | 1222.13 | 322.00 | 810.44 | 343.69 | 828.81 |
| s.e.m. | 7.90 | 5.39 | 45.35 | 33.55 | 8.16 | 24.10 | 15.18 | 23.15 |

